# A promoter::luciferase reporter gene imaging toolkit in *Kalanchoë laxiflora* reveals molecular elements responsible for the circadian regulation of Crassulacean acid metabolism (CAM)

**DOI:** 10.1101/2024.12.19.629446

**Authors:** Jessica H. Pritchard, Jade L. Waller, Peter J. D. Gould, Nirja Kadu, Susanna F. Boxall, Louisa V. Dever, Jana Kneřová, Diarmuid O’Maoileidigh, James Hartwell

## Abstract

Crassulacean acid metabolism (CAM) plants perform primary atmospheric CO_2_ fixation at night, with timekeeping by the endogenous circadian clock. Understanding of circadian coordination of CAM remains limited to rhythmic post-translational regulation of phosphoenolpyruvate carboxylase (PPC) by a specific clock-controlled protein kinase, PPCK. Here, candidate promoter regions (∼3000 bp) of CAM-associated genes from *Kalanchoë laxiflora* were coupled to a firefly luciferase reporter and stable transgenic lines of both *K. laxiflora* and C_3_ *Arabidopsis thaliana* were generated. In *K. laxiflora,* the CAM-associated *GLUCOSE 6-PHOSPHATE/PHOSPHATE TRANSLOCATOR2* promoter (*KlGPT2p*) generated robust circadian rhythms of luciferase luminescence in constant conditions, with peak activity in leaf pair 3, where CAM-associated nocturnal CO_2_ fixation initiated during leaf development. *KlGPT2p::LUC+* did not drive rhythms of luminescence in *A. thaliana* and the *KlPPCK1* promoter produced no LUC+ signal in either species. Furthermore, the *CHLOROPHYLL A/B BINDING PROTEIN2* promoter (*KlCAB2p*), a clock-controlled promoter that drives a gene involved in light-reactions of photosynthesis, drove robust rhythms in both *K. laxiflora* and *A. thaliana*. *KlCAB2p* circadian period changed during leaf development in *K. laxiflora,* revealing differing control by the core-clock during development. *KlCAB2p* peak activity shifted to dawn in *A. thaliana* relative to a dusk phased peak in CAM leaves of *K. laxiflora*, highlighting differences in the timing of outputs from the core clock between species. These findings establish a robust *PROMOTER::LUC+* reporter system in a CAM plant and highlight divergent timing driving clock controlled promoters between species, and period lengthening with leaf age in *Kalanchoë*.

**One-sentence Summary:** Robust circadian rhythms of firefly luciferase in the Crassulacean acid metabolism (CAM) model species *Kalanchoë laxiflora* were driven by both CAM and non-CAM gene promoters.

## INTRODUCTION

The metabolic adaptation of photosynthetic CO_2_ fixation known as Crassulacean acid metabolism (CAM) requires fine-tuned temporal control of metabolism to avoid futile cycling (Hartwell, 2006; Boxall *et al*., 2017). In contrast to plants using C_3_ and C_4_ photosynthesis, which open their stomatal pores for CO_2_ assimilation in the light period, an inversion of stomatal opening and shift of primary atmospheric CO_2_ assimilation to the dark period allows CAM plants to increase water-use efficiency (WUE) by reducing evapotranspiration (Cushman and Bohnert, 1999; Borland *et al*., 2009). This makes CAM a potential tool for achieving sustainable global food security in the face of climate change (Borland *et al.,* 2009). In particular, the forward engineering of a drought-inducible CAM system into existing crop varieties could adapt them to the extreme drought and high temperature events that are becoming more frequent due to the changing climate (Borland *et al.,* 2014).

CAM has evolved from C_3_ photosynthesis at least 66 times independently (Gilman *et al.,* 2023). It serves as a metabolic addition that concentrates CO_2_ around RuBisCO, the ancestral primary carboxylase. During CAM evolution, the timing of metabolic and physiological processes associated with CAM was reprogrammed through changes to the signal transduction cascade(s) that connect the core circadian clock to CAM enzymes and metabolite transporters (Hartwell, 2006). Previous work to identify components responsible for the synchronization of the 24-h cycle of CAM in response to the endogenous clock has established phosphoenolpyruvate carboxylase kinase *(PPCK1)* in the model CAM plant *Kalanchoë fedtschenkoi* as a key circadian regulator of the pathway (Nimmo *et al*., 1987; Carter *et al.,* 1991; Hartwell *et al.,* 1999; Boxall *et al*., 2017). PPCK phosphorylates PPC in the dark period, which reduces its sensitivity to feedback inhibition by malate (Nimmo *et al*., 1984). When *PPCK1* was silenced using RNA interference (RNAi) in transgenic lines of *K. fedtschenkoi*, there was a 66% reduction in nocturnal primary CO_2_ fixation. The disruption of CAM in the absence of clock-controlled *KfPPCK1* led to downregulation and/or perturbation of regulation of several core circadian clock genes, suggesting crosstalk between CAM-associated processes and the core clock mechanism (Boxall *et al.,* 2017). Similar evidence for crosstalk between the metabolic steps of CAM and the core circadian clock was also reported in other transgenic RNAi lines of *Kalanchoë* that lacked key CAM pathway enzymes and lost their ability to operate the CAM cycle, instead reverting to relying on the C_3_ cycle (Dever *et al.,* 2015; Boxall *et al.,* 2020). However, the mechanism that connects *PPCK1* to the core clock so that it is produced *de novo* each night remains unknown (Hartwell *et al*., 1996; Hartwell *et al*., 1999). Gene regulatory network analysis has been applied to *K. fedtschenkoi* to predict candidate circadian clock regulators of CAM, with a specific focus on transcription factors that may bind to and regulate the promoters of core CAM pathway genes (Moseley *et al.,* 2021). In addition, the candidate upstream promoter regions of candidate CAM-associated genes were reported to be enriched for clock-controlled promoter motifs in the CAM crop *Ananas comosus* (pineapple) (Ming *et al.,* 2015). However, none of the numerous published predictions from such *in silico* studies have been tested experimentally in transgenic CAM plants. Furthermore, despite this progress, almost no work has been done to understand the role of *cis*-elements in controlling CAM gene expression.

Transcription factor (TF) mediated timing of gene expression via interactions with conserved motif sequences on promoters of clock-controlled genes (CCGs) is understood most thoroughly from studies in the model C_3_ plant *Arabidopsis thaliana* (Mockler *et al*., 2007). The mechanisms, signaling pathways and individual TFs that regulate specific CAM genes and optimise their timing are not well understood in CAM species, although it has become increasingly clear that the core circadian oscillator components are highly conserved across photosynthetic eukaryotes (Corellou *et al.,* 2009; Lai *et al*., 2020). The core clock genes and their patterns of co-regulation are also broadly conserved in CAM species where this has been studied (Boxall *et al*., 2005; Hartwell, 2006). Other mechanisms have been reported for circadian modulation of gene expression including via post-translational means including protein turnover, calcium signaling, protein phosphorylation and chromatin remodeling (Sugano *et al.,* 1998; Perales and Mas, 2007; Cha *et al*., 2017; Ruiz *et al*., 2018; Uehara *et al.,* 2019).

Firefly luciferase (LUC)-mediated bioluminescence via heterologous expression of the *LUC* gene has been widely adopted as a molecular-genetic reporter by plant scientists. It has been leveraged to elucidate many different aspects of plant molecular biology, including regulation of promoter activity, as a marker gene for successful transformation and as a biosensor. One of the earliest uses of firefly luciferase as a reporter gene in plants coupled *LUC* to the upstream promoter region of the *A. thaliana CHLOROPHYLL A/B BINDING PROTEIN2* (*AtCAB2*) gene and revealed that the activity of this promoter was under circadian clock control (Millar *et al*., 1992a). Coupling the *AtCAB2* promoter upstream of the *LUC* coding sequence, Millar *et al*. (1992a) demonstrated that *AtCAB2* was under transcriptional rather than post-transcriptional or post-translational regulation by the endogenous circadian clock. Since that first seminal report, the use of the *AtCAB2::LUC* reporter in circadian research has led to the discovery of many of the key components of the plant core circadian clock. Random mutagenesis of the *AtCAB2::LUC+* reporter line of *A. thaliana* with ethyl methane sulphonate allowed screening for circadian clock-associated phenotypes via high-throughput imaging for perturbed rhythms of luciferase activity, and such forward genetic screening has led to the identification the major clock component and pseudo response regulator (*PRR*) gene, *TIMING OF CAB2 EXPRESSION1* (*TOC1*), as well as core clock genes *LUX ARRYTHMO* (*LUX*), *EARLY BIRD* (*EBI*) and *TIME FOR COFFEE* (*TIC*) (Millar *et al*., 1995; Strayer *et al.,* 2000; Hazen *et al.,* 2005; Ding *et al*., 2007; Ashelford *et al*., 2011). Many other core-clock gene promoters and candidate clock-controlled promoters of genes that have been discovered to display circadian rhythms of steady-state transcript abundance have been engineered to drive *LUC* in stable transgenic lines. Examples include the promoters of core clock genes *TOC1*, *CIRCADIAN CLOCK ASSOCIATED 1* (*CCA1*), *LHY*, *GIGANTEA* (*GI*), the *EARLY FLOWERING3* (*ELF3*) and *ELF4*, and several *PRR* genes, among many others (Doyle et al., 2002; Nakamichi et al., 2004; Salome and McClung, 2005; Locke et al., 2006; Palagyi et al., 2010; Greenwood et al., 2019; Nimmo and Laird, 2021). Despite being a keystone for understanding plant molecular-circadian systems, a *CAB2* promoter region has not been leveraged for circadian studies in any CAM species and the use of *PROMOTER::LUC* constructs to study circadian biology in other plant species, beyond *A. thaliana*, has been limited to date.

Currently, the sole model system for transgenic studies of CAM are species within the genus *Kalanchoë*, especially *K. fedtschenkoi* and *K. laxiflora* (Dever *et al.,* 2015; Hartwell *et al*., 2016; Boxall *et al.,* 2017; Boxall *et al*., 2020; Lefoulon *et al.,* 2020; Ceusters *et al*., 2021; Hurtado-Castano *et al*., 2023). These species have publicly available genomes and transcriptomes (Yang *et al.,* 2017). Importantly to affirm findings from these data, stable transgenic lines can be generated via tissue culture using *Agrobacterium tumefaciens*, and a diploid accession of *K. laxiflora* that can set viable seed has more recently been identified. This further enhances the amenability of this system as a molecular-genetic model for both CAM and other novel adaptations of plant biology found in the genus *Kalanchoë* (Dever *et al*., 2015; Hartwell *et al*., 2016; Wang *et al.,* 2019).

For the current work, promoter regions of CAM-associated clock-controlled genes (CCCGs) and non-CAM CCGs from *K. laxiflora* were cloned upstream of the *LUC+* reporter and used to generate stable transgenic lines of both CAM *K. laxiflora* and C_3_ *A. thaliana*. *KfPPCK1* has previously been identified as a CCCG that oscillates robustly at the level of its transcript abundance in constant conditions (LL), and its transcript oscillations were blocked by inhibitors of transcription, translation and general protein kinase and calcium/ calmodulin signaling pathway inhibitors (Hartwell et al., 1999; Hartwell et al., 2002). However, the putative upstream promoter region of *PPCK1* has not been studied experimentally for a CAM species, and so the *K. laxiflora PPCK1* promoter (*KlPPCK1p*) was a key target for the *PROMOTER::LUC+* transgenic reporter lines developed here to study the circadian regulation of CAM-associated genes. The second CCCG chosen here was *GLUCOSE 6-PHOSPHATE: PHOSPHATE TRANSLOCATOR2* (*GPT2*), the coding region of which encodes a glucose 6-phosphate:phosphate translocator that is localised to the plastid inner envelope (Kammerer *et al.,* 1998). In the C_3_ species *A. thaliana,* GPT has been shown to have roles in sugar signaling during early seedling development, sucrose-induced leaf growth, and responses to microbial volatiles (Dyson *et al*., 2014; Van Dingenen *et al.,* 2016; Gamez-Arcas *et al.,* 2022). The regulation of *AtGPT2* is linked to light responses via *REDOX RESPONSIVE TRANSCRIPTION FACTOR1* (*RRTF1*), and its response to sucrose is regulated by the transcription factor *AtMYB56* (Jeong *et al*., 2018; Weise *et al.,* 2019). Though there have been advances in understanding the functions of GPT2 in leaves of C_3_ *A. thaliana*, understanding of its induction coincident with CAM is more limited. Haüsler *et al*. (2000) reported that GPT activity increased coincident with CAM-induction in the facultative CAM species *Mesembryanthemum crystallinum* (Common Iceplant). They proposed dual roles for GPT supporting both nocturnal export of G6P from chloroplasts to feed glycolysis and thus phosophoenolpyruvate (PEP) provision for primary CO_2_ fixation, and the uptake of G6P, derived from the recycling of pyruvate generated during malate decarboxylation in the light, to feed starch regeneration in the chloroplast (Hausler *et al*., 2000). *GPT2* transcript abundance was induced coincident with CAM in response to either drought or salt stress, peaked in the light period, and oscillated with a robust circadian rhythm under LL conditions in *M. crystallinum* (Hausler *et al*., 2000; Kore-eda *et al*., 2005; Cushman *et al.,* 2008). Furthermore, the promoter of *McGPT2* was shown to have higher activity in CAM-induced leaves of *M. crystallinum*, supporting the conclusion that the reported transcript level regulation was due to increased transcription driven by the *McGPT2* promoter (Azad *et al.,* 2013). In wild-type *K. laxiflora,* transcripts of *KlGPT2* oscillated robustly under LL free-running conditions in CAM-performing mature leaves (Boxall *et al.,* 2020). Again, in the study silencing *KlPPC1*, leading to a complete loss of dark period atmospheric CO_2_ fixation associated with CAM, *KlGPT2* was also downregulated, emphasising the link between *GPT2* and CAM (Boxall *et al.,* 2020). Whether the activity of the promoter of *GPT2* is responsible specifically for controlling daily, temporal variation in the CAM-associated activity of the protein, in transporting G6P into and/ or out of the chloroplasts, is yet to be determined.

A novel diploid accession of *K. laxiflora* obtained from the living collection at the University of Oxford Botanic Gardens (OBG) was used here to establish a robust luciferase imaging system for studying CCCG promoter activity and investigate links between the core-clock and CAM-associated gene regulation. *K. laxiflora* provided a good system for comparing CAM activity as the youngest leaves flanking the shoot apical meristem, leaf pair 1 (LP1), perform “C_3_-like” CO_2_ fixation, with all net atmospheric CO_2_ assimilation occurring during the light period via RuBisCO, whereas mature LP6 perform strong CAM, with the majority of 24-h atmospheric CO_2_ fixation occurring in the dark period via PPC (Boxall *et al.,* 2020). *PROMOTER::LUC+ (p::LUC+)* fusion gene constructs were generated and used to generate stable transgenic lines expressing *KlCAB2p::LUC+* as a positive control that established a novel *in planta* luciferase imaging system. Furthermore, stable lines expressing a reporter construct for the promoter region of the CCCG *KlGPT2* (*KlGPT2p::LUC+*) led to confirmation that this CAM gene is regulated by the circadian clock at the level of its promoter activity. Comparison of the luciferase bioluminescence rhythmicity driven by the non-CAM *KlCAB2p,* and the CAM-associated promoter *KlGPT2p,* revealed a possible link for *KlGPT2* with the development of CAM as leaves mature in *K. laxiflora* between LP1 and LP6. By contrast, *KlCAB2* promoter activity displayed its highest amplitude rhythmicity in the youngest leaf pairs, consistent with the higher level of light-period CO_2_ fixation associated with C_3_ photosynthesis in the youngest leaves. Detailed characterisation of the circadian rhythms of LUC+ activity driven by *KlCAB2p* in different leaf ages of *K. laxiflora* revealed a changing circadian period as leaves matured. Unexpectedly, the CCCG *KlPPCK1p* was unable to drive any detectable LUC+ activity in multiple independent stable transgenic lines of *K. laxiflora.* Finally, each *K. laxiflora p::LUC+* construct was used to generate stable lines of *A. thaliana* to determine whether the core-clock of this C_3_ species could regulate non-native promoter regions and investigate levels of conservation of promoter regulation between a CAM species and a C_3_ species. When expressed in *A. thaliana*, the timing of peak *KlCAB2p* activity shifted by up to 12-h compared to CAM leaves. Furthermore, bioluminescence activity of the *KlCAB2::LUC+* construct in *A. thaliana* displayed age-dependent circadian period changes, although with shorter periods than those measured in the transgenic lines of *K. laxiflora.* Taken together, these results revealed that the ∼3000 bp *KlCAB2p* possessed conserved promoter motifs that led to it being regulated both temporally and developmentally by endogenous circadian clock associated transcription factors in C_3_ *A. thaliana*. Most importantly though, the results identified fundamental differences in the timing of signaling from the core-clock in C_3_ *A. thaliana* as compared to the CAM leaves of *K. laxiflora*.

## RESULTS

### *KlCAB2p* drove robust rhythms of LUC+ activity in *K. laxiflora*

An experimental pipeline was established to allow the study of *p::LUC+* reporter gene constructs in the model CAM species *K. laxiflora* (Figure 1). In particular, experiments were designed to establish whether or not the promoter regions of CAM-associated genes that are known to be under circadian clock control at the level of their steady-state transcript abundance were responsible for observed transcript rhythms. Both the control CCG *KlCAB2p* and known CCCGs in *K. laxiflora* (Boxall *et al*., 2020) were selected for promoter cloning and analysis in stable transgenic lines of *K. laxiflora* and *A. thaliana*. Binary vector pPCV-LUC+ reported previously (Toth *et al*., 2001) was adapted by introducing the Gateway^TM^ (GW) conversion cassette in the multiple cloning site upstream of the *LUC+* reporter gene coding sequence. This generated the binary vector *pPCV_LUC+_GW* (GenBank Accession Number PQ770919), which was used for the assembly of the *p::LUC+* constructs for the selected *K. laxiflora* genes. Each candidate gene promoter was defined as approximately 3000 bp upstream of the putative ATG start codon and cloned into *pPCV_LUC+_GW* using Gateway^TM^ recombination such that each candidate promoter region was placed immediately upstream of the *LUC+* coding sequence. A *K. laxiflora* diploid accession OBG was stably transformed via tissue culture using methods adapted from those described previously (Dever *et al*., 2015; Boxall *et al*., 2020). Multiple independent transgenic lines were recovered and mature plants ∼30 cm tall were screened for *p::LUC+* activity using a bioluminescence imaging system based on a CCD camera (Figure 1) (Boxall *et al*., 2020).

**Figure 1.**
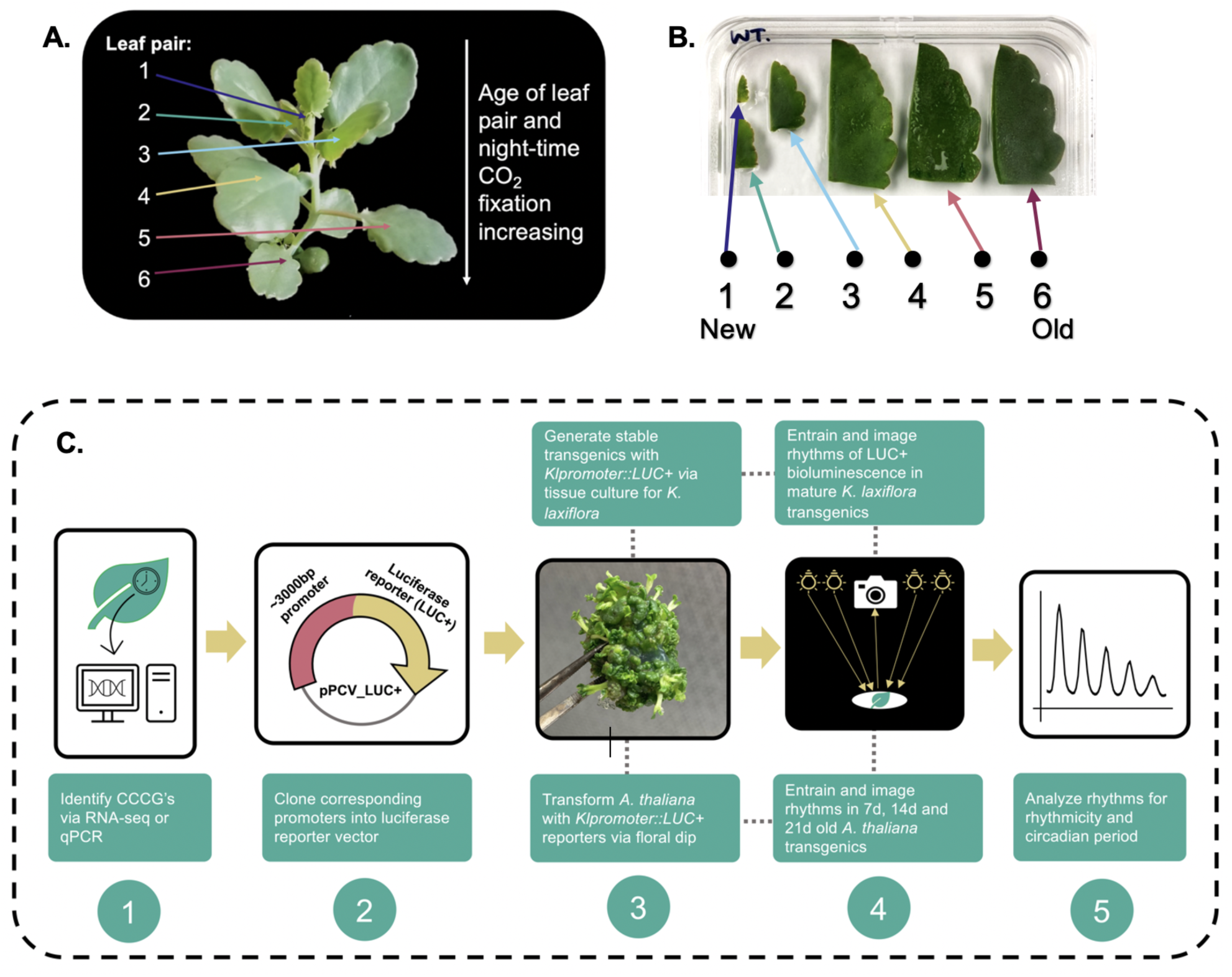
Overview of the pipeline for *K. laxiflora* gene *p::LUC+* reporter construct development and assay in stable transgenic lines of *K. laxiflora* and *A. thaliana*. *K. laxiflora* leaf pairs develop and mature in a sequential manner from the shoot apical meristem down the stem. There is a developmental gradient of CAM-associated CO_2_ fixation as the leaves develop until leaf pair 6 (LP6), where all atmospheric CO_2_ assimilation is occurring in the dark period, and leaves have reached full CAM. Beyond LP6 full CAM continues (e.g. Supplemental Figure 4 in Boxall *et al.,* 2020) **(A)**. Experimental set-up of LP1 to LP6 explants on an agar plate 18-h before luciferase imaging commenced. Half of each leaf was removed leaving the midvein, and the leaf segments were embedded in agar and 50 mM luciferin was sprayed evenly across the plate, saturating each leaf surface **(B).** Pipeline depicting the workflow from identification of candidate CAM-associated clock-controlled CAM genes (CCCGs) (1), through to validation of circadian rhythms of LUC+ driven by corresponding promoters (5) **(C)**.

In *K. laxiflora,* the magnitude and duration of dark period atmospheric CO_2_ assimilation indicative of CAM increases as leaves develop and mature, which in turn enhances the utility of this system for the study of circadian rhythms associated with different levels of CAM (Figure 1A) (Boxall *et al*., 2020). To reduce leaf surface-area and maximise throughput, leaf pairs 1 through 6 (LP1 to LP6, where LP1 is the newly emerging pair of leaves flanking the shoot apical meristem, Figure 1A) were excised from each transgenic line and half of each leaf was cut away, keeping the midvein intact for the half to be imaged. Each leaf segment was embedded in agar to keep it hydrated (Figure 1B). This experimental setup established an *in planta* LUC+ imaging system for *K. laxiflora* that recapitulated LUC+ activity observed for detached intact leaves and leaves on intact plants, but made optimal use of the limited imaging space under the CCD camera’s field of view (Figure 1B).

Due to the extensive previous use of the *AtCAB2p::LUC+* system as a powerful tool to study circadian biology in *A. thaliana*, *KlCAB2p* was targeted initially to establish the equivalent experimental system for monitoring *PROMOTER::LUC+* expression in *K. laxiflora.* Furthermore, the transcript abundance of *Kalanchoë CAB2* was confirmed to oscillate robustly over a 12-h-light/ 12-h-dark cycle from RNA-seq data (Figures 2A & 2C).

**Figure 2.**
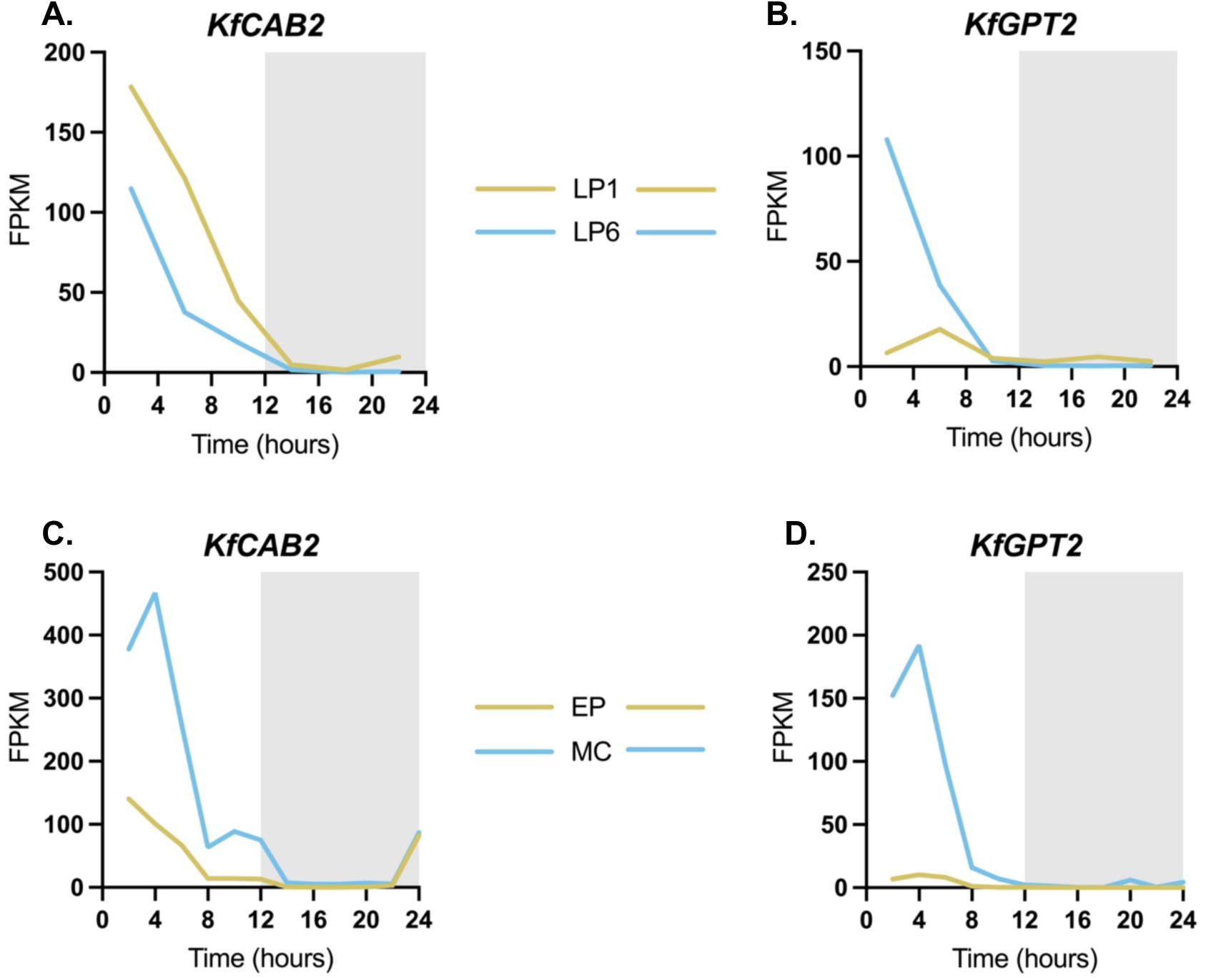
*KfCAB2* and *KfGPT2* transcripts oscillated in abundance over a 12-h-light/ 12-h-dark cycle in both different leaf ages and separated tissue types from mature CAM leaves of *K. laxiflora.* Rhythmic variation over the light/dark cycle of the transcript abundance of *CAB2* **(A & C)** and *GPT2* **(B & D)**. Y-axis values display the Fragments per kilobase of transcript per Million mapped reads (FPKM) generated through RNA-seq analysis. RNA samples from mesophyll cells (MC) and leaf pair 6 (LP6) are in blue whereas samples from epidermal peels (EP) and leaf pair 1 (LP1) are in yellow. Samples were collected from wild-type *K. fedtschenkoi* leaves pre-entrained under 12-h-light/ 12-h-dark cycles and then collected either every 2-h from leaf pairs 6, 7 and 8, with epidermal peels separated from underlying mesophyll cells at each time point **(C & D)** or, in a separate experiment, every 4-h sampling whole LP1 and LP6 **(A & B).**

Stable *K. laxiflora* transgenic lines carrying the *KlCAB2p::LUC+* construct exhibited robust circadian rhythms of bioluminescence for all LPs (Figure 3 and Supplemental Table 1). Reassuringly, the results mirrored previously published results for *AtCAB2p::LUC* in *A. thaliana* (Millar et al., 1992a). The robustness of these free-running rhythms was supported by statistical testing for rhythmicity using Biodare2 (Moore et al., 2014; Zielinski et al., 2014). All LPs were scored highly rhythmic based on a Benjamin-Hochberg P value < 0.005 (Supplemental Table 1). The characteristics of the free-running circadian rhythms of *KlCAB2p::LUC+* varied with leaf age (Figure 3). LP2 had the highest relative bioluminescence peaking at the end of the first subjective light period with a mean relative bioluminescence intensity of 187 (Figures 3A and 3C), revealing that the *KlCAB2* promoter was most active in LP2. However, there was a lot of overlap between the amplitude of the rhythms displayed by leaves of different ages, especially for the youngest pairs of leaves, LP1, LP2 and LP3 (Figures 3A and 3B). For example, the light signal intensity for LP2 was only approximately 1.5-fold more than the next highest in LP1.

**Figure 3.**
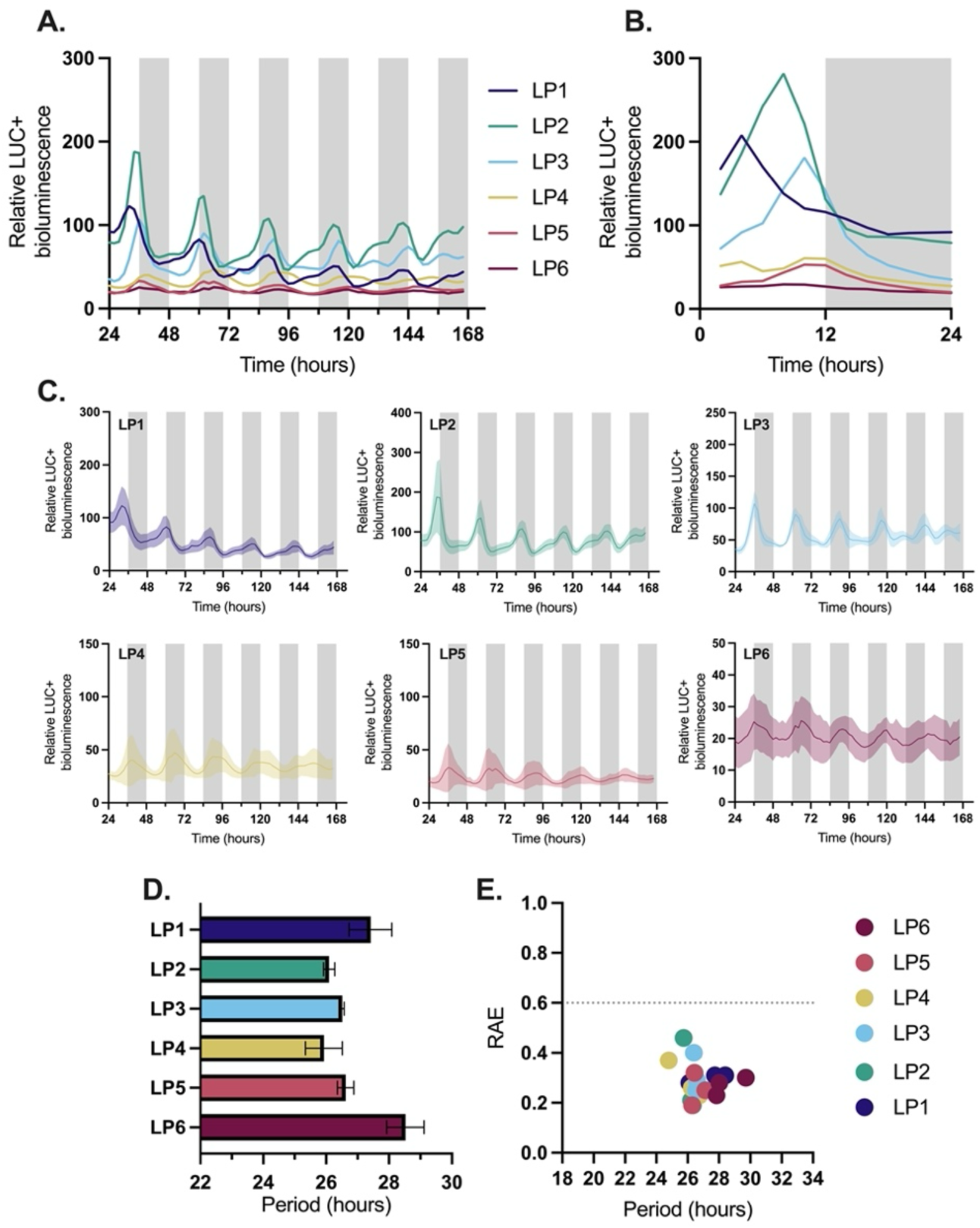
*KlCAB2p::LUC+* drove robust circadian rhythms of luciferase expression in a leaf age dependent manner in *K. laxiflora*. Three biological replicates of a stable transgenic line of *K. laxiflora* diploid expressing *KlCAB2p::LUC+* were imaged in LL for 7-d at 15°C. White and grey bars represent presumptive light and dark respectively. Mean LUC+ expression from for all *K. laxiflora* leaf pairs from 24-h – 166-h **(A)** plotted in ZT. Bioluminescence for the first 24-h in LL for *K. laxiflora* **(B).** 24-h to 166-h of *KlCAB2p::LUC+* expression in *K. laxiflora* for each individual leaf pair. The bold line represents the mean, and shaded area represents SEM. The y-axis scales vary **(C.).** Period and RAE calculated in LL for data from 24 - 96 h for *K. laxiflora* and *A. thaliana* using FFT-NLS. Bars represent SEM. Only replicates that were significantly rhythmic when tested using Biodare2 eJTK and BD2 Classic with a P value < 0.005 after Benjamin-Hochberg correction were included in period calculations. In *K. laxiflora* an ordinary one-way ANOVA showed significant differences between means of period values for all leaf pairs, where P = 0.0158 (P < 0.05) **(D & E).**

For *KlCAB2p::LUC+* the period length of the free-running circadian rhythms increased with leaf age from LP5 to LP6, with an ordinary one-way ANOVA showing a significant difference between the mean period values (Figure 3D). The range of periods measured for this *LUC+* reporter construct were markedly longer than the previously reported free-running period of ∼20 to 21-h for the CAM-associated CO_2_ fixation rhythm under LL conditions in LP6 and whole young plants at the 8- to 10-leaf pairs stage, which were conducted at the same temperature (Supplemental Figure S2 and Boxall et al., 2020).

### The CCCG *KlGPT2p* drove robust LUC+ rhythms with increased amplitude according to leaf developmental stage, but the CCCG *KlPPCK1p* did not generate detectable LUC+

*KlGPT2* transcripts were previously reported to oscillate robustly in older *K. laxiflora* leaves that perform full CAM (Figure 2B & 2D, Boxall *et al.,* 2020). In contrast to the rhythms of *KlCAB2p*, for which rhythm amplitude varied by small increments with leaf age (Figure 3), the mean activity of the *KlGPT2p::LUC+* construct expressed in *K. laxiflora* peaked in LP3, with other LPs, both younger and older, having much lower LUC+ activity levels (Figure 4). For example, after the peak signal in LP3, the next highest bioluminescence signal for *KlGPT2p* was around 3.5-fold lower in LP2 (Figures 4A and 4C). Furthermore, statistical analysis of the rhythms revealed that LP1 were not rhythmic with a high relative amplitude error (RAE) above 0.6 and a Benjamin-Hochberg corrected P value < 0.005 (Supplemental Table 2). All other leaf ages were however rhythmic in at least one replicate based on these tests (Figure 4E & Supplemental Table 2). It was noteworthy that towards the end of the imaging period under LL conditions, there was an increase in the LUC+ activity driven by the *KlGPT2p* in both LP1 and LP2 (Figures 4A and 4C). This response was consistent with the fact that the leaves continued to grow throughout the experimental period under LL conditions, and that after 6-days LP1 were progressing towards the stage that LP2 were at when the time course began, and likewise LP2 were progressing to the LP3 stage (Figure 4). It was furthermore noteworthy that, under LL free-running conditions, *KlGPT2p::LUC+* activity peaked at subjective dusk (Figures 4A, 4B and 4C), which differed from the timing of the peak of the steady-state transcript abundance level of *KlGPT2* at the start of the light period under LD cycles (Boxall *et al.,* 2020).

**Figure 4.**
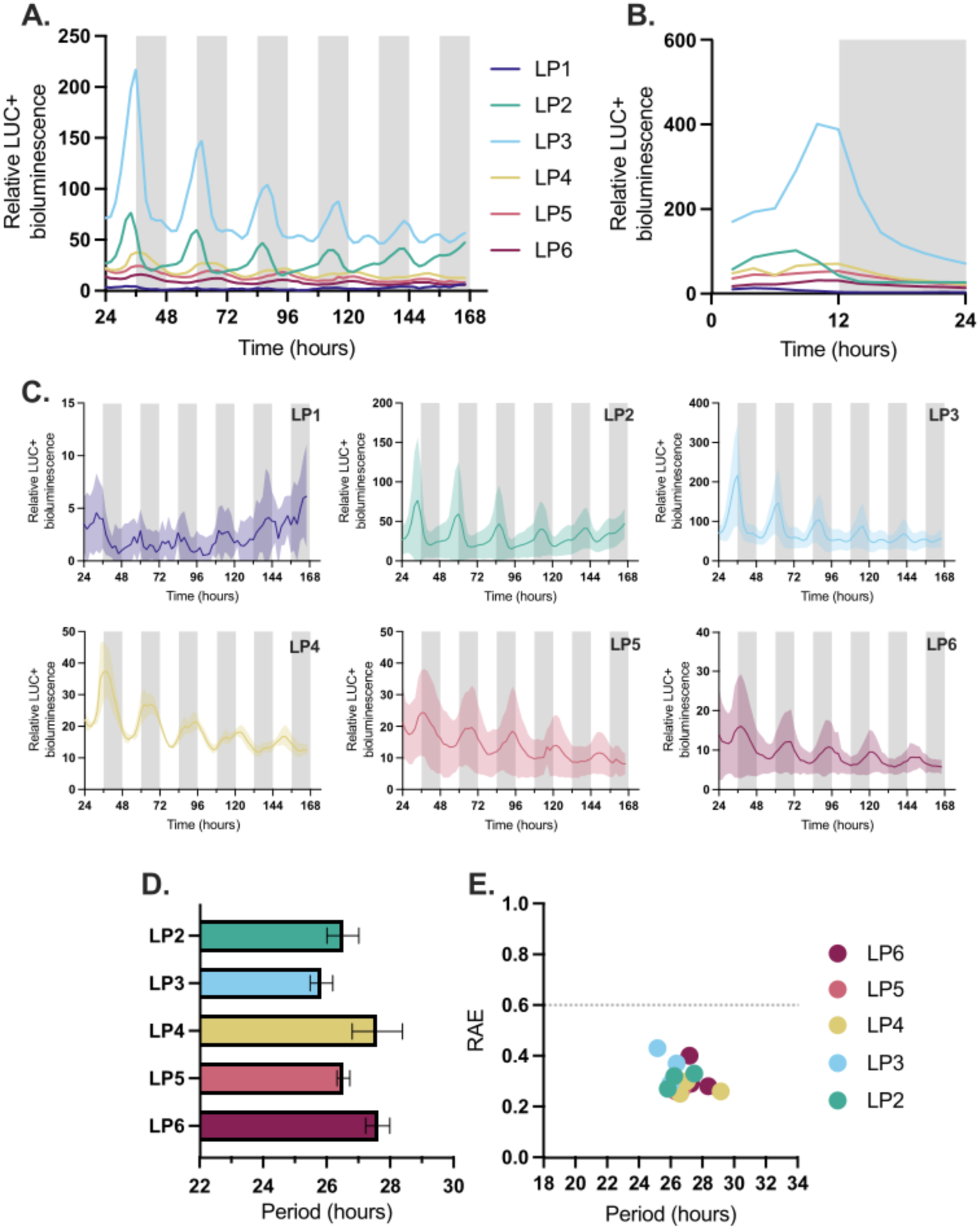
*KlGPT2p* drove robust circadian rhythms in *K. laxiflora* and no detectable LUC+ in *A. thaliana*. Three biological replicates of a stable transgenic line N of *K. laxiflora* diploid OBG expressing *KlGPT2p::LUC+* were imaged under LL conditions at 15°C for 7-d. White and grey bars on graphs represent the subjective light and dark periods, respectively. Mean LUC+ expression from 24-h – 166-h for all leaf pairs plotted in ZT **(A)**. Mean LUC+ expression over first 24-h plotted in ZT **(B).** Twenty-four to 166-h of LUC+ bioluminescence for each leaf pair. The bold line represents the mean, and shaded area represents SEM. Note that the y-axis scales vary **(C).** Period and RAE calculated in LL for data from 24-h – 96-h with FFT-NLS. Bars represent SEM. Only replicates which were significantly rhythmic when tested using Biodare2 eJTK and BD2 Classic with a P value < 0.005 after Benjamin-Hochberg correction were included in period calculations. An ordinary one-way ANOVA showed no significant differences between means of period values for all leaf pairs, where P = 0.8591 (P > 0.05) **(D & E).**

Intriguingly, *KlPPCK1p* did not generate a detectable level of LUC+ luminescence in *K. laxiflora* leaves of any age, with minimal luminescence being detected across the full 7-days in all leaf ages sampled (Supplemental Figure 1). This was despite the robust rhythms of *Kalanchoë PPCK1* transcript abundance that underpin the most thoroughly studied circadian control point for CAM (Hartwell et al., 1999; Boxall et al., 2017).

For both the *KlCAB2p* and the *KlGPT2p* genotypes, the phasing of the first peak of *LUC+* activity during the first 24-h of LL free-running conditions mirrored the trend of the period shifting sequentially as the leaves matured (Figures 3B and 4B). The peak bioluminescence from both *KlCAB2p* and *KlGPT2p* progressed from early into the first subjective light period for LP1 through to the peak being phased at subjective dusk in LP5 and LP6 (Figures 3B and 4B). Taken together, these results for both *KlCAB2p*, which drives a gene associated with the core light-reactions of photosynthesis, and CAM-associated *KlGPT2p,* revealed a change in the properties of the core-clock as leaves age, with a lengthening of circadian period as well as advancing phase to later and later times in the first 12-h subjective light period of each LL experimental run.

### *KlCAB2p::LUC+* drove robust rhythms in *A. thaliana* with a phase shift of peak luminescence timing compared to CAM performing LP5 and LP6 of *K. laxiflora*

The majority of the current understanding of the core circadian clock and clock-controlled gene regulation in plants has been developed from studies in the C_3_ model species *A. thaliana*. The same *KlCAB2p::LUC+*, *KlGPT2p*::*LUC+* and *KlPPCK1p::LUC+* reporter constructs were introduced into *A. thaliana* to elucidate similarities and differences between the promoter activity when expressed in either a CAM or C_3_ species. The resulting stable transgenic lines allowed two key questions to be explored. Firstly, could the *K. laxiflora* promoters be activated in *A. thaliana*, and furthermore, were they coupled to the *A. thaliana* core-clock and able to drive LUC+ expression with a robust circadian rhythm under LL free-running conditions? Secondly, and more importantly in terms of the goal of understanding circadian control of CAM genes, were the temporal characteristics of the *K. laxiflora* promoter reporters in C_3_ *A. thaliana* consistent with the regulation of the same gene promoters in their native environment, or would the promoters generate rhythms with distinct circadian behaviors?

The *KlCAB2* promoter drove robust free-running rhythms of LUC+ activity in 7-day-old *A. thaliana* seedlings, whereas the CAM-associated *KlGPT2p* did not generate a detectable level of LUC+ bioluminescence (Figure 5A). *KlCAB2p* drove relatively low levels of LUC+ expression in the context of the native version, *AtCAB2p*, but interestingly, the distinct promoters drove rhythms with very similar circadian profiles and phasing (Figures 5A and 5B). This included a characteristic peak at the start of the light period. Furthermore, the period lengths measured in these conditions were similar across the two promoters from each species, despite limited sequence similarity or conservation of known circadian clock regulated promoter motifs (Figure 5E).

**Figure 5.**
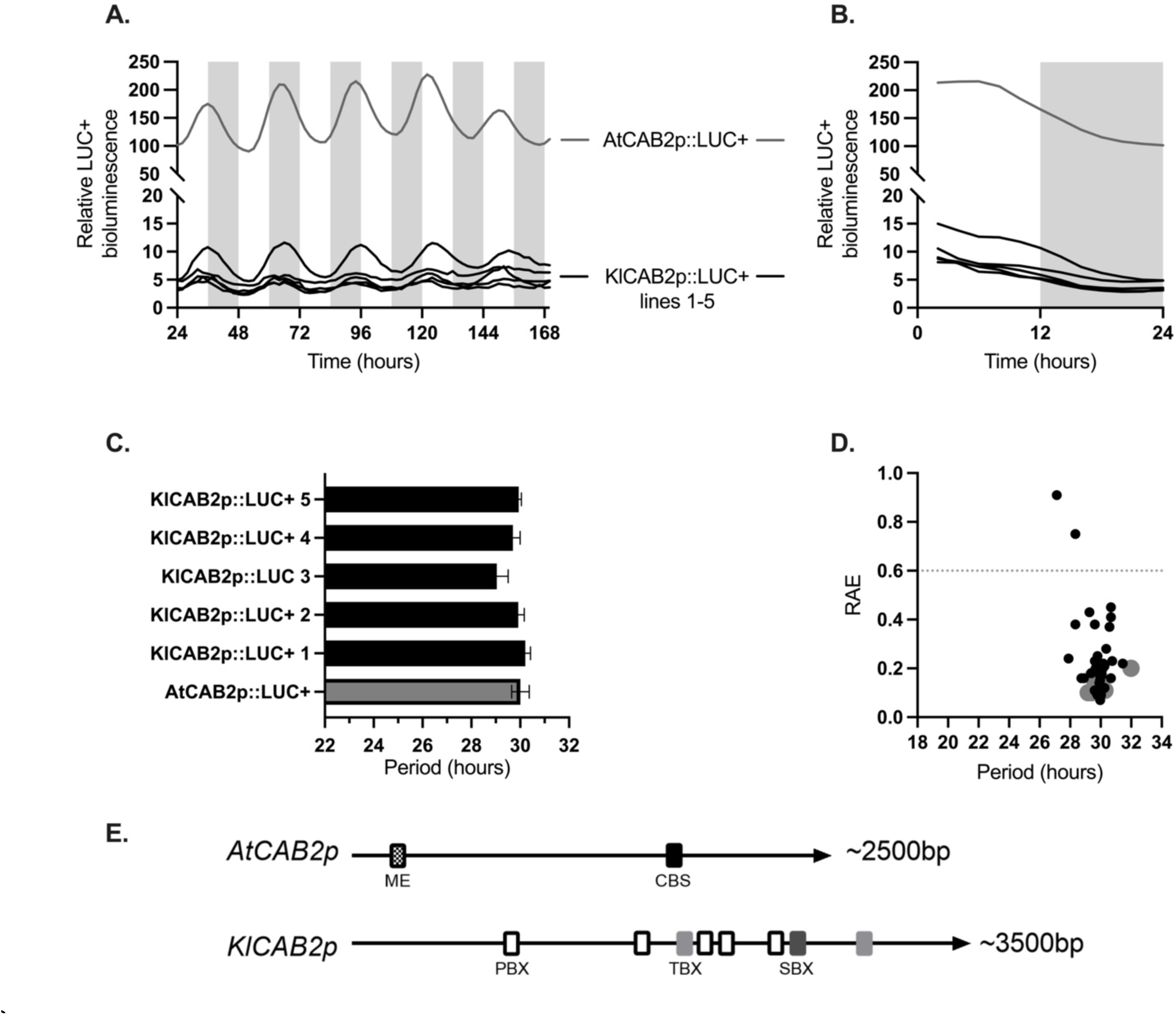
*KlCAB2p* drove robust oscillations with comparable rhythms to *AtCAB2* in *A. thaliana.* Eight replicate wells containing ∼10 seedlings of one line of *AtCAB2p::LUC+* and five independent lines of *A. thaliana* expressing *KlCAB2p::LUC+* were imaged under LL for 7-days at 15°C. White and grey bars on graphs represent the subjective light and dark periods, respectively. Mean LUC+ expression from 24-h – 166-h for all leaf pairs plotted in ZT **(A)**. Mean LUC+ expression over first 24-h plotted in ZT **(B).** Period and RAE calculated in LL for data from 24-h – 96-h with FFT-NLS. Bars represent SEM. Only replicates which were significantly rhythmic when tested using Biodare2 eJTK and BD2 Classic with a P value < 0.005 after Benjamin-Hochberg correction were included in period calculations **(C & D).** Circadian motifs the Morning Element (ME: AACCAC), *CIRCADIAN CLOCK ASSOCIATED 1*-binding site (CBS: AAAAAATCT), protein box (PBX: ATGGGCC), telo-box (TBX: AAACCCT), starch box (SBX: AAGCCC) mapped onto putative promoters for *AtCAB2* and *KlCAB2* **(E)**.

The phasing of *KlCAB2p::LUC+* activity did not match the phase of CAM performing leaves e.g. LP5 and LP6 in *KlCAB2p* in *K. laxiflora* over the first 24h (Figure 5B). This distinction when expressing the same promoter in two species demonstrated that *KlCAB2p* was regulated by the endogenous core circadian clock in *A. thaliana* in the same way as the native *AtCAB2p*. This finding in turn revealed that the *K. laxiflora* core oscillator and its output signaling pathway linking to *KlCAB2* regulation have different circadian properties, and changing properties with leaf development.

To investigate whether the trend of varying period with leaf age measured for *KlCAB2p::LUC+* in *K. laxiflora* leaves (Figure 3D) was also a phenotype for the same reporter construct expressed in C_3_ *A. thaliana,* imaging was performed on individual leaves from 21-d old rosettes of *KlCAB2p::LUC+ A. thaliana*. Leaves were numbered according to a previously published system (Kim *et al.,* 2016), such that the first pair to appear after initial cotyledons were the oldest leaves, numbered 1.1 and 1.2, and the youngest leaf for the 21-d old plants was 5 (note that the second leaf in this pair had not developed) (Figure 5A). *AtCAB2p:LUC*+ was measured according to the same numbering.

Looking at the window from 72-h to 96-h, corresponding to the first day after release into LL, *KlCAB2p::LUC+* activity in *A. thaliana* peaked at subjective dawn (Figure 6C), consistent with the results for 7-d-old plants (Figure 5B). This supports the conclusion that imaging detached leaves did not lead to a fundamental change in the circadian activity of the *KlCAB2p*. As was the case for individual replicates of *K. laxiflora* leaves, *A. thaliana* leaves were discounted from the analysis if they did not pass statistical tests for rhythmicity. *KlCAB2p::LUC+* was able to drive rhythms in detached leaves of *A. thaliana*, but much less robustly than in whole rosettes of plants. However, one replicate from LP1 and 3 replicates for LP4 did generate robust rhythms. Many more of the replicates taken for *AtCAB2p::LUC+* were statistically rhythmic. Firstly, when comparing the two genotypes it was clear that highest relative activity of *KlCAB2p* was in the newest leaves versus *AtCAB2p* where the highest activity was detected in the oldest leaves, LP1 (Figures 6B and 6D). Secondly, the range of period values for rhythmic leaves was significantly longer for *KlCAB2p*, varying from 26.7-h in LP1 to 27.7-h in LP4, than for the native *AtCAB2p*, which showed shorter periods between 23.7-h in LP1 to ∼25.0-h in LP3, and LP4 *AtCAB2p* showed rhythms with periods much closer to the typically expected 24-h. Despite expression in Arabidopsis and a shift in the timing of the peak expression, the *KlCAB2p* period lengths matched closely those of the same promoter expressed in *K. laxiflora* of 26.10-h to 28.52-h.

**Figure 6.**
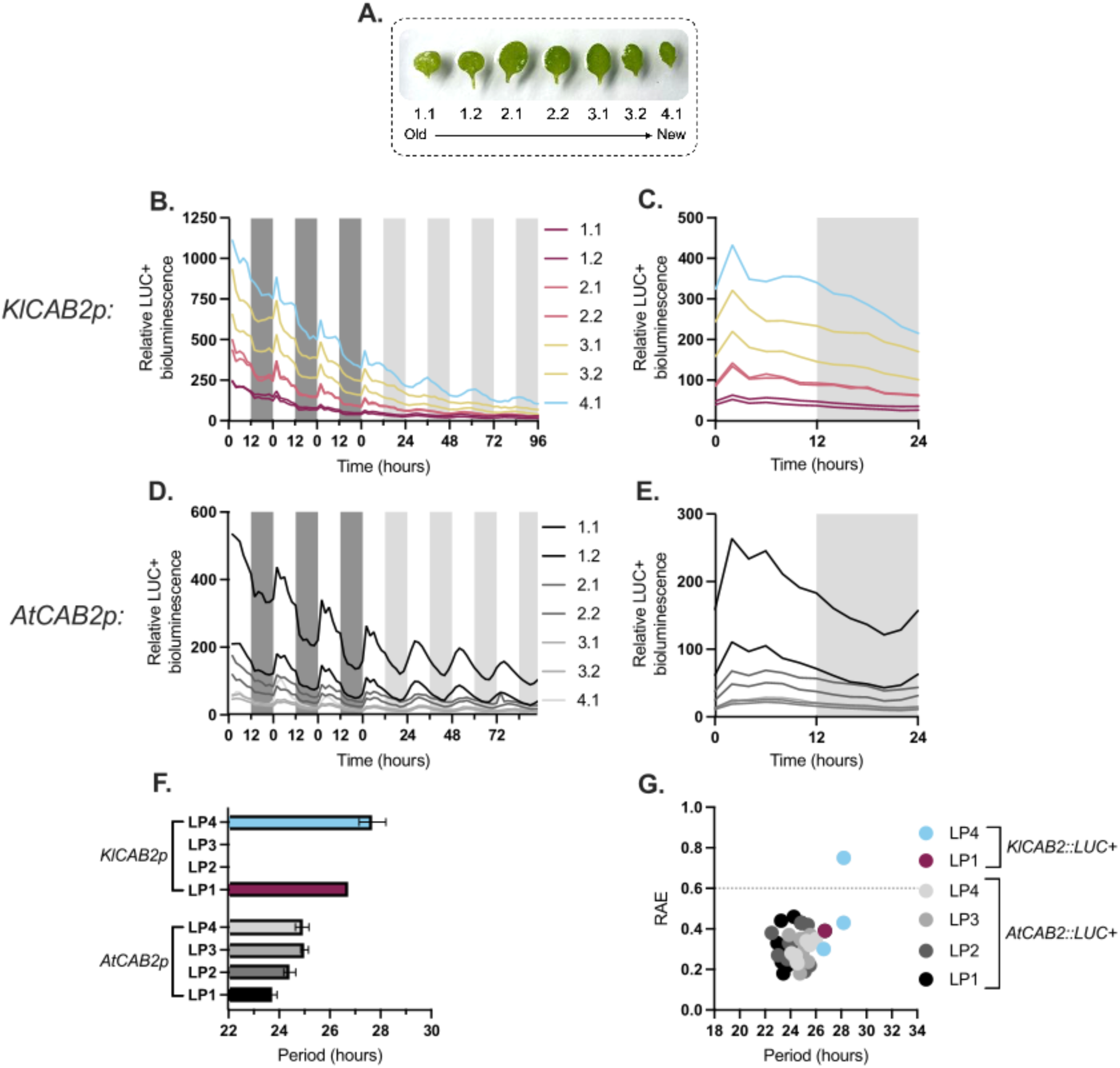
*KlCAB2p* had higher activity in newest detached leaves of *A. thaliana* and *AtCAB2p::LUC+* in the oldest leaves with shorter periods than new leaves. Nine replicates of stable transgenic line no. 4 of *A. thaliana* expressing *KlCAB2p::LUC+* and the line *AtCAB2p::LUC+*, all at 21-days-old were imaged in LD for 3 days and LL for 4 days at 15°C. Dark grey and light grey bars represent true dark and presumptive periods, respectively. Leaf numbering scheme according to leaf age of the 21-d-old *A. thaliana* rosettes **(A)**. Mean LUC+ bioluminescence for each leaf number for entire imaging period plotted in ZT **(B+D)**. LUC+ bioluminescence from 0-h – 24-h plotted in ZT **(C+E)**. Period and RAE calculated in LL with data from 0-h – 72-h with FFT-NLS with linear detrending. Bars represent SEM. Only replicates which were significantly rhythmic when tested using Biodare2 eJTK and BD2 Classic with a P value < 0.005 after Benjamin-Hochberg correction were included in period calculations. An ordinary one-way ANOVA for *AtCAB2p* results showed significant differences between means of period values for all leaf pairs, where P = 0.0056 (P < 0.05). Too many replicates were ruled out from *KlCAB2p* to calculate a P value **(F+G).**

Both *CAB2* promoters drove periods that were longer in younger leaves (Figure 6F). With fewer replicatess passing the statistical test for *KlCAB2p,* it was harder to assess statistically, but there was a clear indication that, for the rhythms scored as robust, the youngest leaves of LP4 displayed longer periods than LP1. There was a strong indication that for the native *AtCAB2p,* periods lengthened by ∼1 h, sequentially between LP1, the oldest leaves, and LP3 / LP4.

## DISCUSSION

Current understanding of plant circadian biology has advanced dramatically thanks to the use of powerful molecular-genetic approaches in the model C_3_ species *A.* thaliana (McClung, 2019). However, this is just one species from the vast diversity of plants, and its regulatory systems will have some degree of uniqueness, especially when compared to species that possess evolved adaptations that are under circadian clock control but are absent from *A. thaliana,* such as CAM (Hartwell, 2006; Hartwell *et al*., 2016). This has contributed to the emergence of other models as amenable molecular-genetic systems for the study of aspects of plant biology that *A. thaliana* doesn’t serve, such as *Kalanchoës* for CAM, *Medicago truncatula* for studying the nitrogen-fixing legume root nodule symbiosis, the genus *Mimulus* as a model for evolutionary genetics, and *Brachypodium* as a model monocot for grass and cereal biology (Hartwell *et al*., 2016; Bandyopadhyay *et al*., 2019; Yuan, 2019; Hasterok *et al*., 2022). In many of these other species, distinct groups of genes are under circadian clock control in comparison to those known to be regulated by the clock in *A. thaliana*. Good examples of this circadian molecular-genetic diversity are the clock-controlled genes underpinning optimised temporal control of complex processes like CAM, and yet the circadian system of CAM species has not been studied in anywhere near as much detail as that in *A. thaliana*. Furthermore, given the interest in engineering a drought-inducible CAM system into C_3_ crop species to enhance their water-use efficiency, it is vital to understand the circadian clock-control of the CAM system (Borland *et al*., 2014; Yang *et al.,* 2015; Lim *et al*., 2019; Schiller and Brautigam, 2021). This is particularly important considering that loss of one circadian regulator of CAM, *PPCK1*, led to a 66% reduction in primary nocturnal CO_2_ fixation via PPC (Boxall *et al*., 2017).

This study began dissecting the molecular-genetic components that allow daily temporal control of CAM using the model species *K. laxiflora* to establish stable transgenic lines expressing firefly luciferase under the control of candidate clock-controlled promoter regions. *K. laxiflora p::LUC+* constructs were also transformed into *A. thaliana* for comparative analysis of their regulation in a C_3_ species. The findings revealed key differences between the activity of the *KlCAB2p* in the two species, which emphasised the need to study circadian molecular-genetics and signaling pathways in the native environment of the promoter of interest.

*KlCAB2p* was able to drive robust rhythms of *LUC+* expression *in planta* in *K. laxiflora* in LL free-running conditions (Figure 3A and 3C). This finding confirmed the potential of leveraging *p::LUC+* reporter gene constructs in transgenic *K. laxiflora* as an easy-to-use model for studying circadian rhythms in live tissues that are performing active CAM. The same degree of rhythm robustness was seen across multiple independent stable transgenic lines, as well as the same trend of promoter expression level being observed across leaf pairs, with LP2 or LP3 having the highest expression of *KlCAB2::LUC+,* but with a lot of overlap between all LPs (Figure 3A.). These findings most probably reflect the fundamental role of the *CAB2* gene in supporting the light-reactions of photosynthesis in all plants, and across the photosynthetic life of a leaf.

Interestingly, when assessing variation of the circadian period with leaf age for *KlCAB2::LUC+* in *K. laxiflora*, the rhythm period lengthened from LP5 to LP6 with LP2/3/4 having periods consistent with LP5 (Figure 3D). The result seen for LP1 (Figure 3D), which displayed a longer free-running period than LP2, could be linked to the fact that this leaf pair was still undergoing photomorphogenesis and building and attaining full photosynthetic competence (Figure 1B); it is noteworthy in this context that LP1 are noticeably paler green than the other leaf pairs. A further intriguing finding here was that the measured period lengths of the *LUC+* reporter constructs were all longer than the circadian period length of the CO_2_ gas exchange measured for LP6 of wild type plants of the same *K. laxiflora* accession (*K. laxiflora* diploid OBG), which varied from 22.45-h – 24.95-h (Supplemental Figure 2). A free-running circadian period of around 24-h is typical for most organisms and the CAM-associated CO_2_ assimilation physiology in *K. laxiflora* OBG was consistent with this, but the rhythmicity characteristics of the *KlCAB2p::LUC+* reporter diverged more widely from 24-h. Some differences could be explained by temperature compensation resulting from the conditions in which plants were imaged, which were used to reflect LL conditions used previously to measure physiology of the closely related *K. laxiflora* tetraploid accession (Boxall *et al*. 2020). However, the differences between intermediate LPs and LP6 were significant. Plasticity of the clock during aging has been recognised previously and has been widely reported in mammalian systems, with period shortening with age in both rats and humans, whereas in insects a lengthening period was reported (Witting *et al*., 1994; Duffy and Czeisler, 2002; Giebultowicz and Long, 2015). For example, in the fruit fly, *Drosophila melanogaster*, period lengthening with age was referred to as ‘age-related decay’, and clock mutants with disturbed rhythms have been shown to have accelerated aging phenotypes, indicating a causal relationship between aging and changed circadian patterns (Giebultowicz and Long, 2015). In plants, circadian period was reported to be shorter by ∼1-h in older leaves relative to younger leaves using the gene *COLD CIRCADIAN RHYTHM AND RNA BINDING (AtCCR2)* fused to luciferase and the *AtCCA1::LUC* reporter as readouts from the clock (Kim *et al.,* 2016). Delayed fluorescence was used to demonstrate different period lengths between leaves and plant ages of wheat (*Triticum aestivum)* and *Brassica napus,* with the conclusion that aging itself was responsible for these changes and that the clock was running faster in young plants before senescence begins (Rees *et al*., 2019; Kim *et al*., 2018). *K. laxiflora* leaves also perform different roles as they age, including increasing levels of nocturnal CO_2_ fixation and malate accumulation associated with CAM (Jones, 1975; Hartwell *et al*., 1999), as well as the transition from sink leaves to source leaves. Such changes in metabolism as leaves develop and mature could explain why genes are required to be initiated at different times in different leaves. Further investigations into whether *KlCAB2p* and other promoters would exhibit a lengthening period, thereby altering the phase relative to dawn and dusk, under entraining light/dark cycles would need to be conducted to reach stronger conclusions. Good candidates for future analysis via *LUC+* reporter constructs would include core-clock gene promoters such as those responsible for the clock-control of *TOC1*, *CCA1* or *GI*, and other CCGs and CCCGs, to determine whether the lengthening of circadian period with leaf age measured here is a genome-wide phenomenon, or whether it is specific to certain subsets of clock-controlled genes such as *KlCAB2p* used here.

*KlGPT2p* drove robust rhythms of LUC+ bioluminescence in all LPs except LP1, and this result was consistent in multiple, independent, stable transgenic lines of *K. laxiflora (*Figure 4A and 4C). A key difference in the temporal expression patterns of LUC+ driven by the *KlGPT2p* compared to *KlCAB2p* in *K. laxiflora* was that one leaf pair, LP3, had 3.5-fold greater peak bioluminescence from 24-h onwards than the next highest expression from LP2. This would indicate a spike in the promoter activity at this specific stage in leaf development, where the CAM-related nocturnal CO_2_ fixation phenotype of LP3 would match that of a weaker CAM species, or intermediate species that performs a mix of CO_2_ being fixed in the day but more at night. Previous reports for the stress-inducible CAM species *M. crystallinum* revealed an increase in *GPT2p* activity upon CAM induction, plus the transcript abundance of *McGPT2* oscillated in both LD and LL conditions in CAM-induced plants (Kore-eda *et al*., 2005; Cushman *et al*., 2008; Azad *et al.,* 2013). Furthermore, Boxall *et al*. (2020) reported that the transcript abundance of *KlGPT2* was reduced and it lost rhythmicity when the CAM pathway ceased to operate due to a lack of *PPC1* in tetraploid *K. laxiflora*.

The results presented here showing that *KlGPT2p* activity was highest in LP3, taken together with the previous understanding that LP3 is where nocturnal CAM-related CO_2_ fixation takes over from CO_2_ fixation via the C_3_ pathway during the light period, suggests that *GPT2p* may be activated developmentally as leaves grow and mature as part of the developmental induction of CAM in *Kalanchoë* (Boxall et al., 2020). In addition, these results for *KlGPT2p* activity and its control during leaf development could also support the proposal that GPT2 plays a reduced role in active CAM as the leaves age through LP4, 5 and 6, wherein the level of CAM rises to the point where almost all atmospheric CO_2_ fixation occurs in the dark. Alternatively, the functional GPT2 protein may remain stable after it is installed in the chloroplast envelopes following the peak of expression in LP3. However, further investigations into the KlGPT2 protein levels, and its transport activity in isolated chloroplasts, measured across the developmental gradient from LP1 to LP6, would be required to understand the balance between changes in promoter activity across leaf ages and the abundance and activity of the encoded protein. These results also emphasise the need to study leaf pairs throughout their development and aging, as previous studies of the transcript abundance of *KlGPT2* sampled only LP6, and important conclusions could be missing without understanding the transcript abundance in intermediate leaves as they develop from using mostly C_3_, light period CO_2_ fixation, to relying on nocturnal CO_2_ assimilation via CAM (Boxall *et al*., 2020). These data also further emphasise the value of using the leaf developmental profile of *K. laxiflora* to study the transition from C_3_ to CAM, as at each collected timepoint multiple developmental stages with a gradual increase in the level of CAM can be sampled and studied comparatively (Hartwell *et al*., 2016).

A commonality between the *KlCAB2p* and *KlGPT2p* activities was the waves in intensity of bioluminescence of LUC+ across the leaves, which can be observed in the timelapse videos (Supplemental Videos). This observation provides further confirmation of the previous finding that the core-clock is actually a complex integration of single-cell autonomous clocks, which are coupled between neighbouring cells (Beck *et al*., 2001; Rascher *et al*., 2001). The data here for waves of LUC+ activity moving across *K. laxiflora* diploid OBG leaves is also similar to waves of clock-controlled expression observed in *A. thaliana* previously, and is very similar to the previously reported waves of circadian delayed fluorescence in mature *K. fedtschenkoi* leaves performing CAM (Gould *et al.,* 2009).

The *KlPPCKp::LUC+* construct used here to generate stable transgenic lines of *K. laxiflora* diploid OBG consisted of ∼2500 bp upstream from the *KlPPCK1* putative ATG start codon. This candidate promoter region was not able to drive detectable LUC+ bioluminescence in *K. laxiflora* or *A. thaliana.* On one hand, this construct therefore served as a good negative control as it demonstrated that there was no autoactivation of *LUC*+ when the transgene is present in *K. laxiflora*. However, the result was counterintuitive to the known regulation of the *PPCK1* transcript, protein abundance and corresponding protein kinase activity, which are all known to peak each dark period in CAM leaves of *Kalanchoë* due to clock-control, leading to phosphorylation of PPC, which in turn makes PPC up to 10-times less sensitive to feedback inhibition by malate (Nimmo *et al.,* 1984; Carter *et al*., 1991; Hartwell *et al*., 1999; Boxall *et al.,* 2017). The negative results for the transgenic lines of *K. laxiflora* diploid carrying the *KlPPCK1p::LUC+* reporter may indicate that the cloned ∼2500 bp putative promoter region does not possess all of the necessary regulatory motifs to drive expression of even low levels of the *LUC*+ reporter. Either key promoter elements are present further than ∼2500 bp upstream, or there may be distal promoter elements at large distances from the *PPCK1* transcribed region. An alternative explanation would be that the *PPCK1* promoter drives transcription constantly and the transcript is periodically turned over such that the steady-state transcript level oscillates every 24-h cycle, but this is not driven by rhythmic fluctuation of the activity of the *PPCK1* promoter. However, this latter explanation can be dismissed because if this was the explanation then the transgenic lines would have displayed a steady and constant level of LUC+ activity that did not vary over time under LL free-running conditions, whereas, in reality, the transgenic lines had no detectable bioluminescence, either rhythmic or continuous. Given the importance of PPCK1 for the optimised circadian control of CAM and the critical need to understand how PPCK1 transcript levels are increased each night to produce the active protein kinase that mediates PPC phosphorylation, further work must explore longer upstream candidate promoter regions from the *PPCK1* gene in *Kalanchoë* in the hope that a longer stretch of the upstream promoter region will replicate the observed regulation of the transcript abundance.

### The *KlCAB2* promoter was able to drive robust LUC+ rhythmicity in *A. thaliana* but CAM-associated *KlGPT2* or *KlPPCK1* promoters did not

When the *K. laxiflora p::LUC+* constructs were transformed stably into C_3_ *A. thaliana* only *KlCAB2p* was able to drive detectable LUC+ bioluminescence that displayed robust circadian rhythms. It was not surprising that the *KlPPCK1* promoter was unable to drive any detectable LUC+ bioluminescence in *A. thaliana*, because the same construct also produced no light in multiple independent stable transgenic lines of *K. laxiflora*. However, the result for the *KlGPT2p::LUC+* construct in *A. thaliana* is more interesting due to the very bright bioluminescence measured for this construct in *K. laxiflora.* Analysis of the sequences and transcript abundance of associated genes of *KlGPT2* and *AtGPT2* revealed key differences that may, at least in part, explain why the molecular-signalling cascades associated with the *A. thaliana* circadian clock were not able to activate and drive transcription downstream of the *KlGPT2p*. Firstly, there are two circadian clock controlled evening element (EE; AAAATATCT) motifs within the candidate upstream region for *AtGPT2p*, which are not present in the 3000bp *KlGPT2p* candidate region used in this work. The EE is necessary for evening specific transcription (Michael and McClung, 2002). Secondly, and perhaps reflecting this lack of EEs in *KlGPT2p*, the transcript levels of the orthologous genes cycle approximately 12-h out of phase with one another, peaking at opposite ends of the light period. *AtGPT2* transcript abundance levels peaked in the first half of the dark period according to an LL RNA-seq time course dataset for wild type *A. thaliana* ecotype Col-0, whereas the *KlGPT2* transcript level peaks at the beginning of the light period in both LD and LL experiments (Boxall *et al*., 2020; Papatheodorou *et al.,* 2020).

### *KlCAB2p* has altered timing of activity when expressed in *A. thaliana* compared to native *K. laxiflora*

*KlCAB2p* had altered timing of activity when expressed in *A. thaliana* compared to native *K. laxiflora* in LP6. Over the first 24-h in 7-d-old *A. thaliana*, *KlCAB2p* generated the highest bioluminescence activity at subjective dawn, and the light level then decreased gradually throughout the subjective light period. By contrast, *KlCAB2p* in LP1 of *K. laxiflora* peaked just after subjective dawn and in all other leaf pairs the peak shifted gradually later in the subjective light period, shifting ultimately to subjective dusk (Figure 3B). This temporal shift in the timing of peak activity of the *KlCAB2* promoter, which was species dependent, emphasised that the promoter coupled to the local clock when expressed in *A. thaliana*, and the *A. thaliana* clock output signaling pathway that controls the endogenous *AtCAB2* promoter activity was also most likely to be regulating the *KlCAB2p* when it was introduced into *A. thaliana*.

To further explore the interactions between plant and leaf development and the circadian clock control of *CAB2*, single leaves from 21-d-old plants were excised from their rosette and numbered from the oldest (LP1) to the newest to appear (LP4). Leaves were arranged on agar to keep them hydrated and individual bioluminescence measurements over time were taken for each leaf from the resulting image stacks (Figure 6A). The results revealed that the rhythmicity profiles of whole plants (Figure 6A) were not lost by detaching leaves, at least for the newest leaves. Furthermore, the same timing with a dawn-phased peak of the bioluminescence signal was evident, and comparable to LP1 expressing *KlCAB2p* in *K. laxiflora,* but shifted compared to older *K. laxiflora* LPs (Figure 6B).

*A. thaliana* has a previously measured endogenous period close to 24 h and the values for *KlCAB2p* here range from a mean of 27.7 h in the newest leaf, L4, to 26.7 h in the oldest leaf with a rhythmic profile, in LP1. Compared to the same promoter in *K. laxiflora* with a range of periods from 26.1 h – 28.52 h in newest to oldest leaves, there was an overlap of the range of mean periods driven by this promoter. A range of periods were also measured for *KlGPT2p* in *K. laxiflora,* and the range was similar to the data for *KlCAB2p* (26.52 h to 27.78 h) revealing that this trend was not specific to *KlCAB2p*. In terms of transcription factor binding sites on the promoters of the two species, *AtCAB2p* had a CCA1-binding site (AAAAAATCT) and a Morning Element (CCACAC) that were absent from *KlCAB2p* (Figure 6E). The presence of these motifs in the *A. thaliana* promoter could explain the dawn-phased activity of *AtCAB2p* versus *KlCAB2p* displaying peak activity closer to dusk in LP2 and older in *K.laxiflora*, although *AtCAB2p* was not transformed into *K. laxiflora*, which would be an interesting experiment for the future.

Whether these periods increased with leaf age in *A. thaliana*, as observed for *KlCAB2p* in *K. laxiflora*, was less clear, as fewer datasets from individual leaves were found to be rhythmic using Biodare. However, there was a significant difference between period lengths of different *A. thaliana* leaf ages, and the youngest leaf (LP4) had the longest period measured, compared to the oldest (LP1). This was also reflected in the data for *AtCAB2p*, where there was a clear trend of period lengthening as new leaves appeared. These results are consistent with those presented by Kim *et al*. (2016) using *A. thaliana*. Importantly, these results raise key questions about the differences between the two species, and the evolution of their circadian biology across the respective leaf developmental profiles. The initiation of CAM in correlation with the process of *K. laxiflora* leaf aging could contribute to an eroding of the circadian rhythm and lengthening period, or the processes may be more unrelated. This only highlights the need for more research into CAM molecular regulation, especially the daily temporal optimisation of CAM by the circadian clock, and transgenic tools such as those demonstrated in the present study will allow this with greater ease.

### Conclusions and future directions

This study demonstrated that the firefly luciferase reporter gene system can be used as a robust, high throughput and reproducible experimental tool to understand the molecular-genetics of the circadian rhythms of CAM-associated genes that underpin the circadian control of CAM in the model species *K. laxiflora.* When comparing this system to C_3_ model *A. thaliana*, we also uncovered fundamental differences in the clock control over specific promoter regions across the two species, with *KlCAB2p* driving rhythms with different characteristics to those it generates in *K. laxiflora* when it was expressed in *A. thaliana.* This opens up some interesting questions regarding the ability of a host species’ circadian system to couple to promoter motifs from another species and control gene activity over time, especially given the long-term goal of engineering mechanisms from a CAM system with optimised temporal control in order to improve the WUE of C_3_ crops. The data here demonstrate that it is unlikely that coding region transgenes driven by their native corresponding promoters will be sufficient to achieve the correct timing of gene expression in the target plant. Instead, there will be a need to understand all of the circadian timing interactions in the plant being improved. Going forward with these investigations, it would be useful to image more *Klp::LUC+* fusions including some of core-clock genes such as *TOC1* or *CCA1/LHY*, and *K. laxiflora* plants containing these constructs could be manipulated with stresses or other environmental stimuli, as has already been explored in some detail through circadian biology studies in *A. thaliana*. Such studies will enhance understanding of how the core-clock is affected by environmental perturbations in other species that use clock-control to achieve temporal optimisation of novel aspects of plant biology that are absent from *A. thaliana*. For instance, plants could be grown under elevated CO_2_ to investigate the response of the circadian system to the increasing global atmospheric CO_2_ levels that are driving climate change. Other reporters should be investigated for compatibility with *Kalanchoë* species, which could improve the ease and power of such investigations, such as an auto-luminescent fungal bioluminescence pathway (FBP), which would remove the need for application of the luciferin substrate and allow for *in situ* imaging of plants (Khakhar *et al*., 2020). Furthermore, other luciferase reporter gene innovations like nanoLUC, which is a more stable reporter protein than LUC, will prove valuable for monitoring circadian regulation of protein abundance through in-frame fusions of candidate circadian clock-controlled proteins, such as *PPCK1*, to the nanoLUC coding sequence (Khakhar *et al*., 2020; Urquiza-García and Millar, 2019)).

It will also be important to study the proteins related to promoters in this study. For instance, in the case of *GPT2*, the findings here emphasise the need to study the activity of the chloroplast envelope transporter that the gene encodes in order to determine whether the protein abundance and activity remains stable in CAM performing leaves after the spike in promoter activity in LP3. Finally, it will be interesting to determine the mechanistic basis for the interaction between the core-clock of *A. thaliana* and the promoter regions from *K. laxiflora* used here. Specifically, exploring motifs or transcription factor binding sites that can be identified within these ∼3000bp regions as being essential for regulatory elements from the clock-output pathway to interact with them and drive the observed rhythm of promoter activity. Understanding such interactions and signaling networks better would ultimately allow for temporal control to be optimised when considering engineering CAM into C_3_ crops.

## METHODS

### Plant materials

The *Kalanchoë laxiflora* plants used for the generation of transgenic lines were originally obtained from the University of Oxford Botanic Gardens (OBG), UK. Ploidy measurements using flow cytometry determined that this accession is diploid with an estimated genome size of 274 Mb, and self-pollination trials established the ability of this line to set viable seed. The seed used here for stable transformation were the progeny of 3 rounds of self-pollination and single seed descent. For seedling growth and regeneration of transgenic plants via tissue culture, seeds were surface sterilised with ethanol and germinated on ½ x Murashige and Skoog (MS) medium with Gamborg’s B-5 vitamins (Duchefa) and 3% (w/v) sucrose (Sigma) containing 0.8% (w/v) Phytoagar (Duchefa). Plates of seedlings were cultured in walk-in growth chambers set to provide 16-h-light/ 8-h-dark at a constant temperature of 22°C. Mature transgenic lines and wild type *K. laxiflora* were grown in compost mix 1:1:1:3 M2 (Multi-Purpose Compost, Levington M2): John Innes (Sinclair’s John Innes No. 2 Compost): Sinclair compost (Sinclair Potting compost): Perlite (Sinclair Perlite Standard) with additional nutrients from Osmocote beads (ICL Osmocote Pro Slow-Release Fertiliser 5-6), which was added according to manufacturer’s instructions. Mature plants were grown in heated research glasshouses with a minimum temperature of 20°C (seasonally dependent) and supplementary lighting from sodium metal halide lamps ensuring a year-round minimum daylength of 16-h-light/ 8-h-dark. Plants were entrained for circadian free-running experiments using Snijders Microclima MC-1000 plant growth cabinets set to 12-h-light (450 µmol photons m^−2^ s^−1^), 25°C, 60% humidity / 12-h-dark, 15°C, 70% humidity.

*Arabidopsis thaliana* plants of ecotype Col-0 were grown at constant 22°C with16-h-light/ 8-h-dark in research glasshouses and plant growth rooms. For selection of transformants, *A. thaliana* seed were surfaced sterilised using 70% ethanol and plated to germinate on ½ x MS medium with Gamborg’s B-5 vitamins, 0.8% (w/v) Phytoagar and 15 µg ml^−1^ hygromycin B (Duchefa).

### Cloning of *KlGPT2p*, *KlCAB2p* and *KlPPCK1p* regions and construction of *p::LUC+* binary vectors

Putative promoter regions of ∼3000bp were selected upstream of the candidate 5’ transcription start site (TSS) of the target gene, and included the 5’UTR up to the predicted ATG start codon. For *KlGPT2p* and *KlPPCK1p*, cloning primers were designed against the tetraploid ‘*Kalanchoë laxiflora* v1.1’ genome available on the JGI Phytozome database, with the accession numbers Kalax.0007s0180.1 (*KlGPT2*) and Kalax.0021s0061.1 (*KlPPCK1*). *KlCAB2p* was amplified and cloned from the diploid *K. laxiflora* OBG accession that was used to generate all the transgenic lines.

Each ∼3000 bp promoter fragment was amplified from wild type *K. laxiflora* genomic DNA using gene specific primers with Gateway^TM^ ready attB recombination sites added as 5’ extensions. These primer extensions allowed the resulting PCR products, amplified from *Kalanchoë* genomic DNA with KOD Hot-Start DNA polymerase, to be recombined into the pDONR201 Gateway^TM^ entry vector using a BP Clonase recombination reaction according to the manufacturer’s instructions (Thermo Fisher Scientific, UK). Each *Kalanchoë* promoter insert in a pDONR201 entry clone was sequenced in full using a commercial Sanger Dideoxy sequencing service (Eurofins) to confirm the correct promoter sequence had been cloned in the correct orientation. Standard Gateway® cloning was carried out to recombine promoter regions into the *pPCV_LUC+_GW* binary vector (GenBank Accession number: PQ770919) (Katzen, 2007). Plasmids were transformed into competent cells of *Agrobacterium tumefaciens* strain GV3101 pMP90RK using electroporation and allowed to recover through growth in super optimal medium with catabolic repressor (S.O.C.) for 3-h at 30°C on an orbital shaker at ∼225 rpm. The cells were pelleted in a microfuge and resuspended in 200 µL of S.O.C. and both 180 μL and 20 μL were plated out on separate LB-agar plates containing rifampicin 50 μg mL^−1^, gentamycin 25 μg mL^−^1, kanamycin 50 μg mL^−1^ and carbenicillin 100 μg mL^−1^ for selection of positively transformed colonies.

### Generation of transgenic lines containing *p::LUC+* reporter gene constructs

#### Stable transformation of *K. laxiflora* diploid accession OBG using *A. tumefaciens* and a tissue culture regeneration system

Diploid *K. laxiflora* tissue from 4- to 6-week-old seedlings that had been germinated and grown in sterile, tissue culture conditions was used for generation of all stable transgenic lines. Roots were removed from plantlets and leaves were cut into small explants of approximately 5 mm wide using a sterile scalpel in a laminar flow bench. Explants were submerged in the appropriate strain of *A. tumefaciens* GV3101 pMP90RK harbouring the desired *pPCV_LUC+_GW* binary vector construct for 1-h and blotted on sterile filter paper before being transferred to co-cultivation plates. The regeneration of callus and plantlets was done as previously described by Dever *et al*. (2015) with the exception of the antibiotics used. Hygromycin selection at working concentration 5 µg mL^−1^ was used to select for successful transformation of plant material through the presence of *pPCV_LUC+_GW p::LUC+* binary vector and Timentin at 300 µg mL^−1^ was used to prevent over-growth of *Agrobacterium* (Wang *et al*., 2019).

Once plantlets with roots were obtained in tissue culture, they were transferred into the soil mix and grown in conditions described above. They were grown first in plugs ∼5 cm wide until they measured ∼10 cm tall and had established a substantial root network. Plants were then transferred to 12 x 12 cm pots and grown to a height of ∼30 cm prior to their entrainment to 12-h-light/ 12-h-dark conditions in preparation for imaging experiments.

#### Floral dip transformation of *A. thaliana*

Flowering *A. thaliana* plants aged 4- to 6-weeks were dipped once in the appropriate liquid culture suspension of *A. tumefaciens* transformed with the desired *pPCV_LUC+_GW p::LUC+* construct. The *A. tumefaciens* cells were grown overnight in 250 mL LB medium. They were then mixed directly with 250 mL 2X concentration infiltration media to give a final concentration of 10% (w/v) sucrose, 0.025% (v/v) Silwett L-77. Post-dip, plants were watered until first seeds dropped and then allowed to dry for ∼3 weeks, at which point the seed was harvested and cleaned (Clough and Bent, 1998).

T0 seeds were selected on 15 µg ml^−1^ Hygromycin according to protocol by Harrison *et al*. (2006). Resulting T1 plants were genotyped using diagnostic PCR amplification of a ∼400bp fragment that spanned the junction region between the 3’ end of each *K. laxiflora* promoter and the 5’ end of the open reading frame of the *LUC+* gene. T2 seeds were collected from positive clones and used in imaging experiments at 7-days, 14-days or 21-days after vernalisation in the cold for at least 48-h.

#### Luciferase imaging of transgenic lines and image analysis

Plants were entrained in 12-h-light/ 12-h-dark for 7-days prior to imaging, with growth chamber conditions as described above. For *K. laxiflora*, leaf pairs 1 through 6 were excised from each plant at the base of their petiole using a scalpel around 18 to 24-h before imaging and the scalpel was used to remove half the leaf and the petiole, but with the leaf half to be imaged retaining the mid-rib. Each leaf explant was then placed on plates containing 0.4% agar with water (pH 5.8). They were sprayed evenly with a solution containing 5 mM luciferin in 0.01% (v/v) Triton X-100 in water, filter sterilised. Plates with leaves were then returned to the growth cabinet under 12-h-light/ 12-h-dark entrainment conditions for the following ∼18h, until the start of the next light period. For each time course imaging experiment of luciferase bioluminescence, plates of pre-entrained, excised leaves were transferred to the imaging cabinet set to a constant light intensity of ∼30 µmoles photons m^−2^ s^−1^ and a constant temperature of 15°C. Illumination was provided by a mix of red and blue LEDs as described previously (Gould *et al*., 2009). The software ‘micromanager’ connected to a programmable Arduino computer was used to control a Retiga LUMO^TM^ CCD camera. Imaging commenced at the time that would have been the start of the 12-h-light period during pre-entrainment. The red and blue LEDs were switched off before each exposure of the camera, which lasted 20 mins. Images were captured every 2-h spanning a period 7-to 10-days under constant light and temperature (LL), circadian free-running conditions.

#### Data analysis of LUC+ imaging measurements and statistical analysis of circadian rhythmicity and period length

Image stacks were analysed in Fiji (Schindelin *et al*., 2012), selecting a small region of the leaf towards the distal end in the image, and measuring the ‘mean gray value’ of pixels as a quantitative measure of the amount of bioluminescence. Background light levels, measured away from the leaves, were quantified in the same way, and subtracted to normalise data across each time course of images.

For circadian analysis and statistics, including rhythmicity and period estimates, Biodare2 (www.biodare2.ed.ac.uk) was used (Moore et al., 2014; Zielinski et al., 2014). Period was calculated using linear detrending and Fast Fourier Transform-Non-linear Least Squares (FFT-NLS), and then compared to spectral resampling (SR) to ensure validity of results. Rhythmicity was also determined using Biodare2. Data would be filtered for non-rhythmic replicates prior to period calculations (Supplemental Tables 1 - 4).

## Supporting information

Supplemental Tables 1 through 4

## ACKNOWLEDGEMENTS

This research was supported in part by the U.S. Department of Energy Office of Science, Genomic Science Program (grant DE-SC0008834) and in part by the Biotechnology and Biological Sciences Research Council (BBSRC grant BB/F009313/1 to J.H.). J.H.P. was supported by a PhD studentship funded by the BBSRC as part of the Newcastle-Liverpool-Durham Doctoral Training Partnership (DTP). The contents of this article are solely the responsibility of the authors and do not necessarily represent the official views of the DOE.

## Supplemental figure and movie legend

**Supplemental Figure S1. The *KlPPCKp::LUC+* construct did not drive rhythms of LUC+ activity in 4 independent transgenic lines of *K. laxiflora*.** Each graph (on individual tabs of the Excel file) shows quantified luminescence data (y-axis) traces for different leaf ages (LP1 through to up to LP12, depending on the independent line and replicate) imaged over either 4-d (graphs for replicates E1_2, E2_2, I2_2 and I4_2) or 7-d (graphs for replicates E1, E2, I2 and I4) under LL at 15°C. The x-axis of every graph shows the cumulative time under LL in hours.

**Supplemental Figure S2: *Kalanchoë laxiflora* accession OBG diploid displayed a robust and persistent circadian rhythm of CAM-associated CO_2_ exchange under constant light and temperature (LL) free-running conditions.** Three clonal mature plants that had been grown in a research glasshouse for 4-months under 16-h-light/ 8-h-dark were pre-entrained to 12-h-light/ 12-h-dark cycles in a Snijders Microclima MC-1000 plant growth cabinet (12-h-light period conditions: 450 µmoles photons m^−2^ s^−1^, 25°C, 70% humidity; 12-h-dark conditions: lights off, 15°C, 70% humidity). Leaf pair 6 (LP6) were detached from the three clonal replicate plants and their petiole placed in distilled water immediately. Pairs of leaves were set up for measurements in the whole plant chambers of a bespoke CIRAS-DC-based, 12-channel, gas-switching infra-red gas analyser system, as described in Dever et al. (2015) and Boxall et al. (2020). Conditions during the experimental run were initially the same 12-h-light/ 12-h-dark used for pre-entrainment for the first 24-h, with the 12-h-dark period denoted by the darker grey background on the graph, and then the leaves were switched to LL condition after the first 24-h of LD. LL conditions were constant light (100 µmoles photons m^−2^ s^−1^) at 15°C and 70% humidity. The three biological replicate plants are plotted in black, red and blue.

**Supplemental Figure 3. Time lapse movies showing LUC+ activity over ∼1 week in multiple replications of LP1 to LP6 of *K. laxiflora* for the genotypes *KlCAB2p::LUC+* (left) and *KlGPT2p::LUC+* (right).** Images were captured every 2-h for ∼ 7-d under LL at 15°C. Video can also be accessed at https://youtu.be/HdpArmDxA6M?si=z_H5GIqJ7U2C1Sjr

