## Supplemental Figure S2 for "A promoter::luciferase reporter gene imaging toolkit in *Kalanchoë laxiflora* reveals molecular elements responsible for the circadian regulation of Crassulacean acid metabolism (CAM)"

### Slide 1
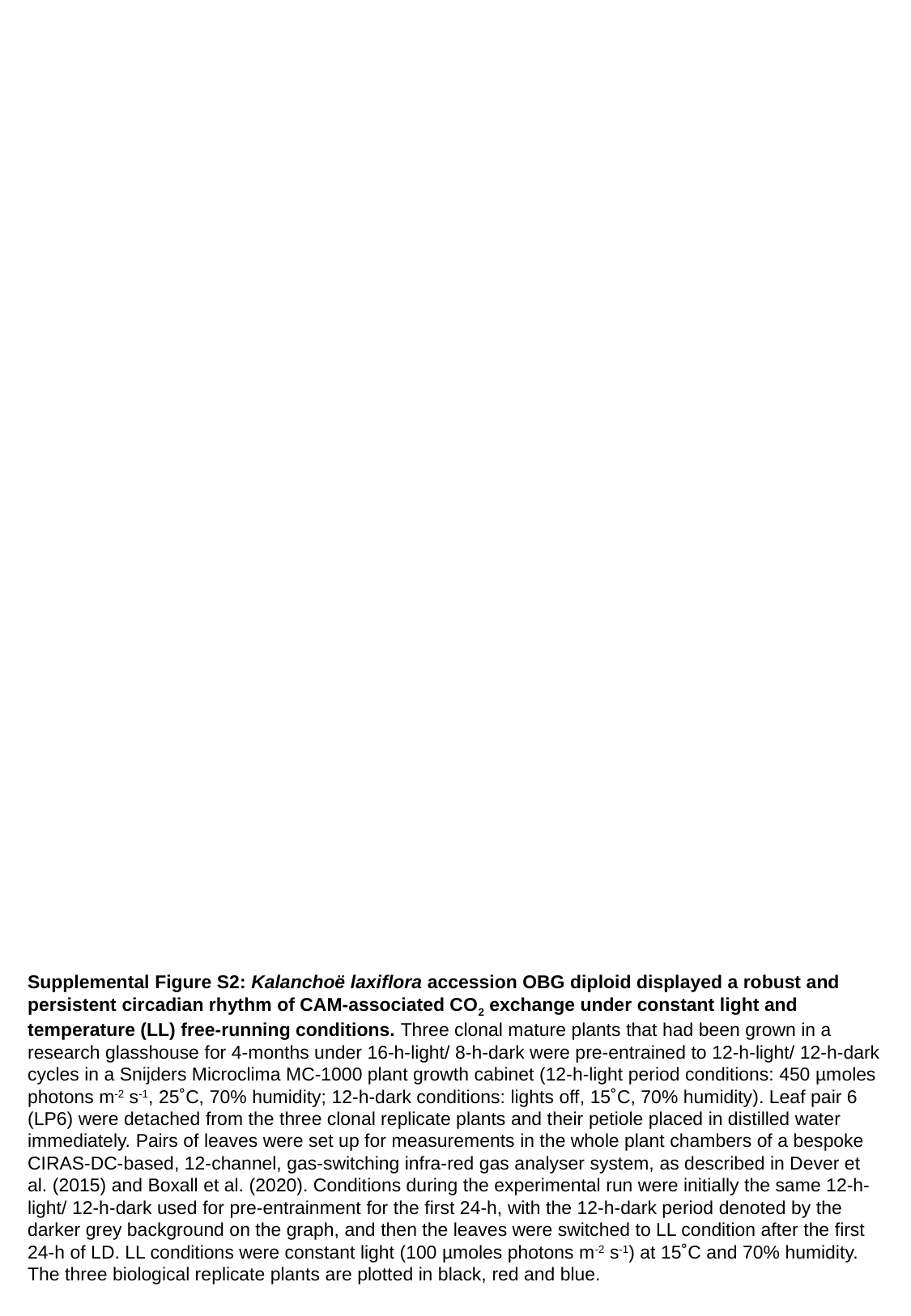

Supplemental Figure S2: Kalanchoë laxiflora accession OBG diploid displayed a robust and persistent circadian rhythm of CAM-associated CO_2_ exchange under constant light and temperature (LL) free-running conditions. Three clonal mature plants that had been grown in a research glasshouse for 4-months under 16-h-light/ 8-h-dark were pre-entrained to 12-h-light/ 12-h-dark cycles in a Snijders Microclima MC-1000 plant growth cabinet (12-h-light period conditions: 450 µmoles photons m^-2^ s^-1^, 25˚C, 70% humidity; 12-h-dark conditions: lights off, 15˚C, 70% humidity). Leaf pair 6 (LP6) were detached from the three clonal replicate plants and their petiole placed in distilled water immediately. Pairs of leaves were set up for measurements in the whole plant chambers of a bespoke CIRAS-DC-based, 12-channel, gas-switching infra-red gas analyser system, as described in Dever et al. (2015) and Boxall et al. (2020). Conditions during the experimental run were initially the same 12-h-light/ 12-h-dark used for pre-entrainment for the first 24-h, with the 12-h-dark period denoted by the darker grey background on the graph, and then the leaves were switched to LL condition after the first 24-h of LD. LL conditions were constant light (100 µmoles photons m^-2^ s^-1^) at 15˚C and 70% humidity. The three biological replicate plants are plotted in black, red and blue.
